# Soil Metatranscriptomes Reveal Phylogenetic Filtering of Candidatus Udaeobacter Across Paired Prairie and Agricultural Sites

**DOI:** 10.64898/2026.08.07.743348

**Authors:** Samuel Lord

## Abstract

Long-term agricultural conversion is known to shift soil microbial diversity and abundance in soils that formerly supported native grassland, but whether these shifts reflect uniform suppression across a bacterial genus or selective filtering of specific evolutionary lineages remains poorly understood. We addressed this question in *Candidatus* Udaeobacter, a globally abundant member of the phylum Verrucomicrobiota and a model oligotrophic soil bacterium. We collected 40 soil samples for RNA-Seq metatranscriptome analysis across three paired native prairie and long-term agricultural sites in Missouri and mapped transcriptional recruitment against a taxonomically curated consensus reference built from 36 concordant NCBI and GTDB *Candidatus* Udaeobacter genome assemblies. Total transcriptional recruitment to *Candidatus* Udaeobacter was nearly eleven-fold higher in prairie soils, and recruitment composition remained significantly distinct between land uses even after normalizing for this difference, indicating that land use reshapes which lineages remain active rather than uniformly reducing activity across the genus. This land use-associated recruitment showed strong phylogenetic signal, with closely related genomes exhibiting similar responses to land use. Genome architecture tracked this pattern and prairie-enriched lineages carried consistently smaller genomes and expressed a larger share of their coding capacity than agriculture-enriched lineages. These results show that environmental selection in Candidatus Udaeobacter operates below the genus level. Combining curated reference genomes with metatranscriptomic recruitment offers a scalable framework for resolving lineage-level ecological responses in other abundant, poorly characterized microbial taxa.

## Introduction

The tallgrass prairie is characterized by a canopy dominated by C4 grasses and an exceptionally diverse forb community, creating one of the most biologically complex terrestrial ecosystems in North America. These millennial-scale natural communities, developed over thousands of years and predating the agricultural conversion that now dominates the region, are some of the most diverse assemblages of plants on earth [1]. At their historical extent, tallgrass prairies covered much of the central United States, and the deep, organic-rich soils beneath them accumulated carbon and structure over long timescales under perennial grassland vegetation [2]. Modern agricultural systems have altered these soils substantially [3], and prairie restoration studies show that while management can improve soil structure and carbon inputs, it does not fully recreate the original prairie soil legacy [4,5]. The aboveground vegetation represents only part of the system, because a large fraction of prairie biomass is belowground in roots and rhizomes, and the diversity and functioning of the soil microbial community are a major component of the ecosystem. Prairie (grassland) systems without a history of soil disturbance house a microbial community shaped over millennia by interactions with edaphic formative processes, plant community dynamics, and abiotic interfaces. This community did not assemble quickly, and it does not appear to reassemble quickly either, once disturbed [6]. The result is a foundational biological structure that drives nutrient cycling and carbon turnover across tall grass prairie ecosystems [7], one that reflects not just the plants growing above it, but the full depth of time the system has had to organize itself.

Within those microbial communities, known constituents of the Verrucomicrobiota phylum are considered ubiquitous and active members of the soil community [8]. Some are known for being genomically streamlined and closely associated with C4 grass communities [9]. This association is not incidental. Root exudate chemistry and rhizosphere carbon inputs differ substantially between C4 grasslands and the row-crop systems that typically replace them [10–12]. A streamlined, oligotrophic organism like *Candidatus* Udaeobacter (Ca. Udaeobacter from here on), may be particularly sensitive to that shift due to the physical, chemical, and biological changes associated with production agriculture, which effectively alter the ecological landscape of the rhizosphere [13,14]. Because Verrucomicrobiota abundance can decline sharply, in some systems by as much as 80%, following conversion to agriculture [15], we developed a hypothesis to ascertain what fragments of the Ca. Udaeobacter genus persist post-conversion, and whether those fragments are consolidated to a different phylogenetic portion of the known Ca. Udaeobacter reference genomes (cross referenced NCBI and GTDB databases), comparatively. Determining whether surviving populations represent a true subset of the ancestral prairie community, or a functionally distinct lineage that happened to persist, has direct implications for how we interpret Ca. Udaeobacter’s role, and by extension the role of oligotrophic carbon specialists more broadly, in degraded versus intact soil systems. This makes tallgrass prairie a useful system for asking not just whether Ca. Udaeobacter survives conversion, but what conversion does to this soil microbial lineage.

We built a consensus reference from taxonomically concordant Ca. Udaeobacter genomes, resolving discrepancies between NCBI and GTDB classification before mapping transcript recruitment across paired prairie and agricultural sites in Missouri, each representing a direct, or proximity adjacent comparison between remnant prairie and the agricultural land immediately bordering it or in close proximity to it, to determine whether these activity differences were concentrated within specific lineages or distributed evenly across the genus. Genome size also emerged as a relevant factor, with clear differences between land uses in the genomic architecture of recruited lineages, suggesting that genomic streamlining in Ca. Udaeobacter is not simply a fixed background trait [9], but something actively shaped by land use. These comparisons are drawn from three site pairs within a single region, so these findings speak most directly to the specific land-use transition captured by this design, and broader generalization across the full range of tallgrass prairie conversion will require testing in other geographic and management contexts.

## Materials and Methods

### Site selection

Sampling targeted three remnant tallgrass prairie sites and four adjacent agricultural fields spanning a range of historic prairie geography across Missouri. Each remnant was paired with one or more nearby agricultural fields sharing the same soil series and comparable topographic position, a design intended to isolate land use as the primary driver of observed differences while holding edaphic and climatic conditions roughly constant between members of a pair.

Two of the three remnants, PennSylvania (Penn) and Golden prairies, sit in southwestern Missouri, a region that retains the largest concentration of intact prairie remaining in the state despite less than 1% of Missouri’s historic prairie extent surviving today. Both sites were chosen for soils and plant composition, which is still representative of undisturbed tallgrass prairie. Each is separated from its paired agricultural field by only a county road, with both members of the pair situated on matching soil series and topographic position; these fields are referred to throughout as Penn ag and Golden ag.

The third remnant, Tucker Prairie, lies in central Missouri and is one of the largest surviving examples of claypan, glacial till prairie in the state, distinguished by loess-derived till soils and a floristic composition that has never been altered by tillage. Tucker Prairie is paired with two agricultural fields, Ag A and Ag B, both within miles of the remnant and situated on the same heavy claypan till soils that historically supported this prairie type.

### Study design and metatranscriptomic dataset

Forty soil samples were collected from paired sites for metatranscriptomic analysis. Metatranscriptomes were used to evaluate transcriptional recruitment to Ca. Udaeobacter across native prairie and long-term row crop agricultural systems indicative of the region. The dataset consisted of 17 prairie metatranscriptomes collected from the Golden, Penn, and Tucker prairie sites and 23 agricultural metatranscriptomes collected from the adjacent Golden ag, Penn ag, Ag A, and Ag B cropping systems. These locations represented three independent prairie and agriculture comparisons: Golden prairie versus Golden ag, Penn prairie versus Penn ag, and Tucker prairie compared with the nearby Ag A and Ag B agricultural fields. This paired sampling design allowed us to evaluate broad differences between native prairie and agricultural management while accounting for geographic pairing during downstream statistical analyses. This study focused exclusively on transcriptional recruitment to curated Ca. Udaeobacter reference genomes. Rather than examining the complete soil microbial community, this analysis was designed to determine which evolutionary lineages of Ca. Udaeobacter were transcriptionally active across contrasting land-use histories and whether recruitment patterns were associated with land use type.

### Sample preparation and sequencing

Soil samples were collected from the paired prairie and agricultural sites described above during 2023 and stored at −80°C until processing. Total RNA was extracted using the RNeasy PowerMax Soil Pro Kit (Qiagen) following manufacturer protocols, and residual DNA was removed using the TURBO DNA-free Kit (Thermo Fisher Scientific). RNA quantity was assessed using a Qubit Fluorometer, and RNA integrity was evaluated using an Agilent Fragment Analyzer prior to library preparation. Samples meeting quality requirements underwent ribosomal RNA depletion and library preparation using the Illumina Stranded Total RNA Prep with Ribo-Zero workflow. Sequencing was performed at the Argonne National Laboratory Environmental Sample Preparation and Sequencing Facility (ESPSF) on an Illumina NextSeq 2000 platform, generating paired-end 150 bp reads. Raw reads underwent quality filtering prior to alignment against the consensus Udaeobacter reference described below.

### Construction of the consensus Ca. Udaeobacter reference set

Publicly available bacterial genome assemblies labeled as *Candidatus* Udaeobacter were retrieved from the NCBI Assembly database using the NCBI Datasets command-line interface using TaxID 1921511 [16]. This initial search returned 90 genome assemblies whose organism names contained Ca. Udaeobacter. Because organism names associated with public genome assemblies do not always reflect current genome-based taxonomy, accession numbers for all 90 assemblies were compared against the Genome Taxonomy Database [17], GTDB release r232. Assemblies were retained only when the NCBI accession was present in GTDB and GTDB assigned that accession to the genus Ca. Udaeobacter. Assemblies whose NCBI organism names contained Ca. Udaeobacter but were classified elsewhere by GTDB were excluded from downstream analyses. Applying this accession-level concordance criterion reduced the initial collection of 90 NCBI-labeled assemblies to 36 taxonomically concordant genomes. These 36 NCBI and GTDB concordant assemblies constituted the consensus Ca. Udaeobacter reference collection used for all subsequent phylogenomic reconstruction, transcript recruitment, differential recruitment, and phylogenetic analyses.

### Transcript recruitment

The 36 taxonomically concordant Ca. Udaeobacter genomes were combined into a consensus reference consisting of 9,564 contigs. Paired-end reads from each of the 40 soil metatranscriptomes were aligned independently to this reference using Bowtie2 v2.4.2 with the “very-sensitive preset” [18]. Resulting alignments were sorted and indexed with SAMtools v1.20. Genome-level recruitment was quantified using samtools idxstats [19], and contig-level counts were aggregated to their corresponding genome accession. This produced a recruitment matrix containing 1,440 genome-by-sample observations representing 36 consensus genomes across 40 metatranscriptomes. Alignment to the consensus reference yielded a total of 1,668,793 mapped reads. Of those mapped reads, 652,459 reads were assigned to predicted coding sequences and retained for gene-level expression analyses. Predicted coding sequences were functionally annotated by assigning KEGG Orthology (KO) identifiers using KofamScan with the default adaptive score threshold [20].

### Taxonomic validation of recruited reads

To independently evaluate the taxonomic specificity of recruitment to the consensus reference, recruited reads were queried against the GTDB r232 representative protein database. Protein-level matches were linked to their corresponding GTDB genome accessions and taxonomic assignments. For each recruited read, the highest-scoring protein matches were evaluated at the genus level, and reads for which Ca. Udaeobacter represented the highest-scoring or co-highest-scoring assignment were classified as independently supported. Taxonomic support was summarized for each metatranscriptome as the proportion of GTDB-classifiable recruited reads receiving Ca. Udaeobacter support. Differences in taxonomic support between prairie and agricultural samples were evaluated using a Wilcoxon rank-sum test.

### Phylogenomic reconstruction

Phylogenomic relationships among the 36-consensus Ca. Udaeobacter genomes were reconstructed using GTDB-Tk v2.6.1 [21] with the GTDB r232 reference database. A concatenated bacterial marker gene alignment was generated for all genomes and used to infer a maximum likelihood phylogeny in IQ-TREE v3.1.3 [22]. The optimal amino acid substitution model was selected using “ModelFinder” according to the Bayesian Information Criterion (BIC), and branch support was estimated by using 1,000 bootstrap replicates. The final phylogeny included all 36 genomes and exhibited strong overall branch support. Of the 33 internal branches with bootstrap values, 17 (51.5%) received support ≥95 and 28 (84.8%) received support ≥70. Bootstrap values ranged from 52 to 100 (median = 95; mean = 88.4), indicating that most relationships among the consensus genomes were well resolved. This phylogeny served as the evolutionary framework for all subsequent recruitment and phylogenetic signal analyses.

### Statistical analyses

All statistical analyses were performed in R v4.6.1 [23]. Differences in total Ca. Udaeobacter recruitment between prairie and agricultural soils were evaluated using Wilcoxon rank-sum tests. Differential recruitment among the 36 consensus genomes was assessed using DESeq2 [24] with the design formula ∼ Pair + Habitat (“land use” for purposes outside of coding), where agriculture was treated as the reference level. Resulting *P*-values were adjusted using the Benjamini-Hochberg procedure to control for false positive discovery [25], and genomes with an adjusted *P* ≤ 0.05 and an absolute log2 fold change of at least ≥0.5 were considered differentially recruited.

Genome recruitment profiles were Hellinger transformed [26] prior to calculation of Euclidean distances for multivariate analyses. Differences in community composition were evaluated using Euclidean distances, principal coordinates analysis (PCoA), and PERMANOVA while accounting for spatial pair as implemented in the vegan package [27]. Homogeneity of multivariate dispersion was evaluated using PERMDISP [28].

Phylogenetic signal was evaluated by mapping genome-specific DESeq2 log2 fold changes in transcript recruitment between prairie and agricultural soils onto the maximum-likelihood phylogeny. Positive log2 fold changes represented greater relative recruitment in prairie, whereas negative values represented greater relative recruitment in agriculture. Blomberg’s *K* [29,30] and Pagel’s λ [31] were estimated using the “phytools” package in R, with significance assessed by 9,999 randomizations for Blomberg’s *K* and a likelihood-ratio test against λ = 0 for Pagel’s λ [29,32,33]. Ancestral land use states were reconstructed using maximum likelihood under alternative Markov models of character evolution implemented in phytools, with model selection based on Akaike’s Information Criterion (AIC). Relationships between genome recruitment breadth and total recruitment were evaluated using Spearman rank correlations.

Because larger reference genomes provide more sequence available for transcript recruitment, genome size was evaluated as a potential source of bias in the recruitment analyses. Genome size, contig number, N50, largest contig, and GC content were calculated for each of the 36 consensus genomes directly from the assembled reference sequences (Table S1). Spearman rank correlations were then used to test whether genome size was associated with total transcript recruitment, recruitment breadth, or land use-associated recruitment. Land use-associated recruitment was represented by genome-specific DESeq2 log2 fold changes, allowing these comparisons to reflect the relative contribution of individual genomes after accounting for spatial pair and differences in overall recruitment among samples. These analyses tested whether genome architecture was associated with lineage-specific responses to land use rather than differences in total Ca. Udaeobacter transcriptional recruitment.

## Results

### Differential recruitment among consensus genomes

Differential recruitment analysis identified substantial land use-specific recruitment across the consensus Ca. Udaeobacter collection. Of the 36 reference genomes, 17 were classified as prairie-enriched and ten as agriculture-enriched, whereas the remaining nine genomes did not meet the combined statistical significance and effect-size criteria after accounting for spatial pair (Figure 1). The 36 consensus genomes ranged from 1.39 to 4.05 Mb and contained between 51 and 589 contigs (Table S1). N50 values ranged from 5.12 to 97.09 kb, while GC content remained highly conserved across the collection, varying from 53.65% to 55.87%. The broad range of genome sizes and assembly completeness reflects the diverse origins of the publicly available assemblies, whereas the narrow GC range is consistent with the taxonomic coherence of the consensus reference collection. Prairie-enriched genomes exhibited log2 fold changes ranging from 1.05 to 3.69, indicating substantially greater transcriptional recruitment in native prairie soils. In contrast, agriculture-enriched genomes exhibited log2 fold changes ranging from −0.52 to −2.20. The most prairie-associated genome (GCA_036269155.1) exhibited a log2 fold change of 3.69, whereas the strongest agriculture-associated genome (GCA_035694525.1) exhibited a log2 fold change of −2.20. Across predicted coding sequences, 597,308 reads were assigned in prairie metatranscriptomes compared with 55,151 in agricultural metatranscriptomes, representing an approximately 10.8 fold difference in gene-assigned transcriptional recruitment, demonstrating that Ca. Udaeobacter was significantly more transcriptionally active under native prairie conditions. To determine whether long-term land use also altered the composition of transcriptionally active lineages independent of this overwhelming difference in recruitment, subsequent analyses were performed on Hellinger-normalized recruitment profiles.

**Figure 1.**
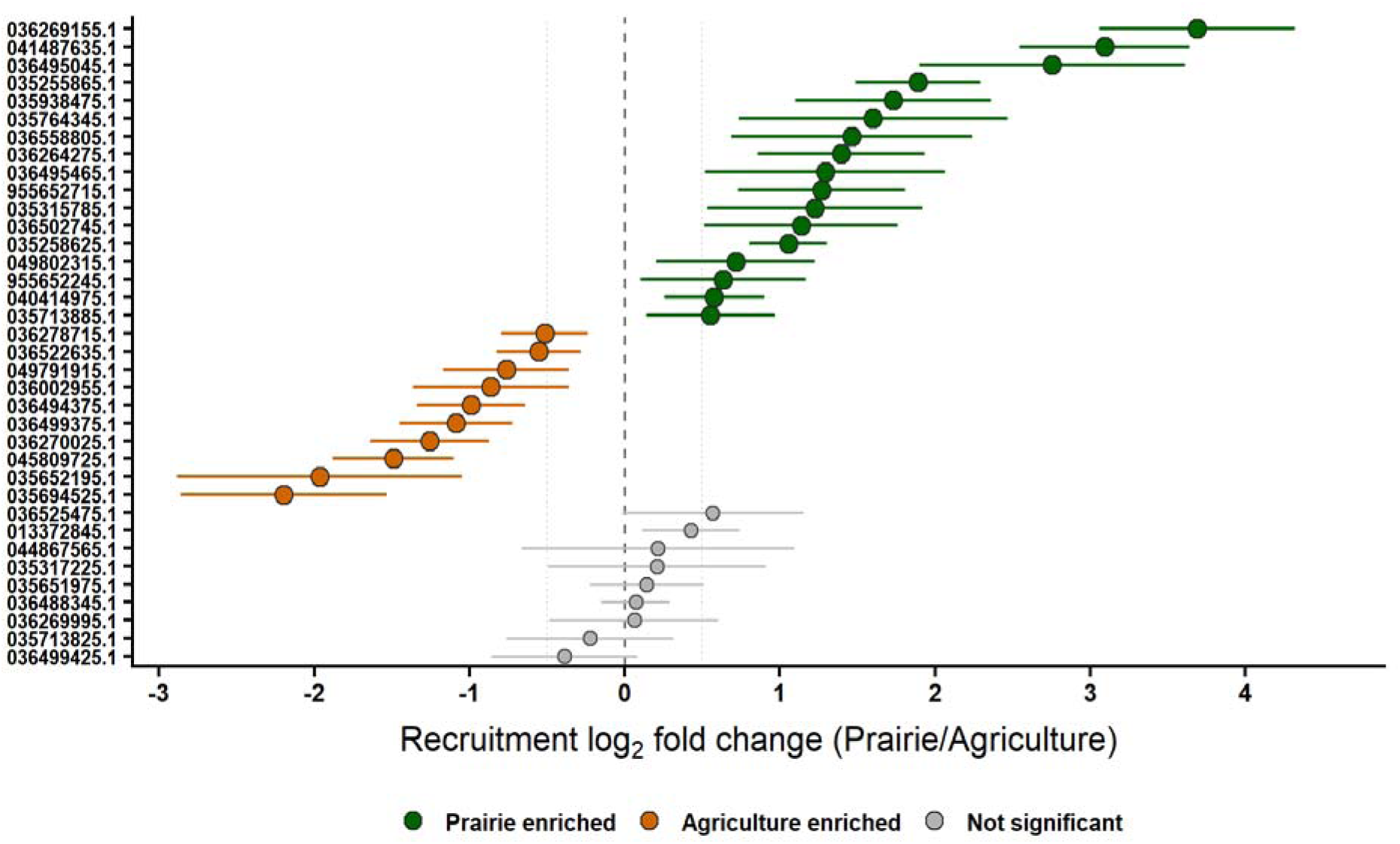
Differential transcriptional recruitment of 36 consensus Ca. Udaeobacter genomes between native prairie and agricultural soils. Differential recruitment analysis based on DESeq2 showing log2 fold changes in transcript recruitment for each consensus genome after accounting for spatial pair. Positive values indicate greater transcriptional recruitment in prairie soils, whereas negative values indicate greater recruitment in agricultural soils. The Y-axis notes the accession numbers of all 36 reference genomes. Points represent estimated log2 fold changes and horizontal bars denote 95% confidence intervals. Green points indicate genomes significantly enriched in prairie soils, orange points indicate genomes significantly enriched in agricultural soils, and gray points represent genomes that did not meet the significance criteria (adjusted *P* < 0.05 and |log2 fold change| ≥ 0.5). Seventeen genomes were significantly prairie-enriched, ten were significantly agriculture-enriched, and nine showed no significant land use preference. **Alt text:** Forest plot of log2 fold change in transcript recruitment for 36 Candidatus Udaeobacter genomes, comparing prairie to agricultural soils. Genomes are ordered from most agriculture-enriched to most prairie-enriched, with points colored green for significant prairie enrichment, orange for significant agriculture enrichment, and gray for no significant difference; horizontal bars show 95% confidence intervals.

Recruitment was also evaluated for paired-read support to assess whether alignments to the Ca. Udaeobacter reference were primarily driven by isolated sequence matches. In a representative prairie metatranscriptome, 84.1% of recruited reads had their paired mate also map to the consensus reference, with 58.6% occurring as properly paired alignments. Only 15.9% of recruited reads occurred as singletons. Thus, most recruitment was supported by both reads originating from the same sequenced fragment rather than by isolated read alignments to the Ca. Udaeobacter reference.

Taxonomic specificity of reference recruitment was further evaluated by independently querying recruited reads against the GTDB r232 representative protein database. Across all samples, 918,301 recruited reads produced classifiable protein-level matches, of which 366,132 (39.9%) had Ca. Udaeobacter as the highest-scoring or co-highest-scoring genus-level assignment. Taxonomic support differed strongly by land use, with a median of 51.0% of classifiable recruited reads independently supported as Ca. Udaeobacter in prairie samples compared with 15.0% in agricultural samples (Mann–Whitney U = 30, P < 0.0001). Lower taxonomic specificity was concentrated in agricultural rather than prairie samples, indicating that non-specific recruitment was not responsible for the observed prairie enrichment of Ca. Udaeobacter.

### Compositional differentiation independent of total recruitment

The large difference in total Ca. Udaeobacter recruitment between prairie and agricultural soils was accompanied by a significant shift in the composition of the recruited genomes. After normalizing recruitment within each sample and applying a Hellinger transformation, land use explained 39.5% of the variation in genome composition (PERMANOVA, R² = 0.395, F = 26.02, *P* < 0.001) after accounting for spatial pair. In contrast, spatial pair explained only 6.5% of the variation and was not significant (R² = 0.065, *P* = 0.084), indicating that land use was the dominant factor structuring Ca. Udaeobacter recruitment profiles (Figure 2). Multivariate dispersion also differed significantly between land uses (PERMDISP, F = 5.31, *P* = 0.025), with agricultural samples exhibiting greater dispersion than prairie samples, indicating that recruitment profiles were more variable among agricultural soils. Despite this difference in dispersion, the strong land use effect observed by PERMANOVA, together with the differential recruitment analysis, supports the conclusion that prairie and agricultural soils harbor compositionally distinct transcriptionally active Ca. Udaeobacter assemblages. Principal coordinates analysis supported this result, with prairie and agricultural samples forming distinct compositional groups using Hellinger normalization. This separation indicates that the land use effect was not driven solely by the much greater abundance of Ca. Udaeobacter transcripts in prairie soils. This would indicate long-term agricultural management altered the relative contribution of individual genomes within the recruited assemblage.

**Figure 2.**
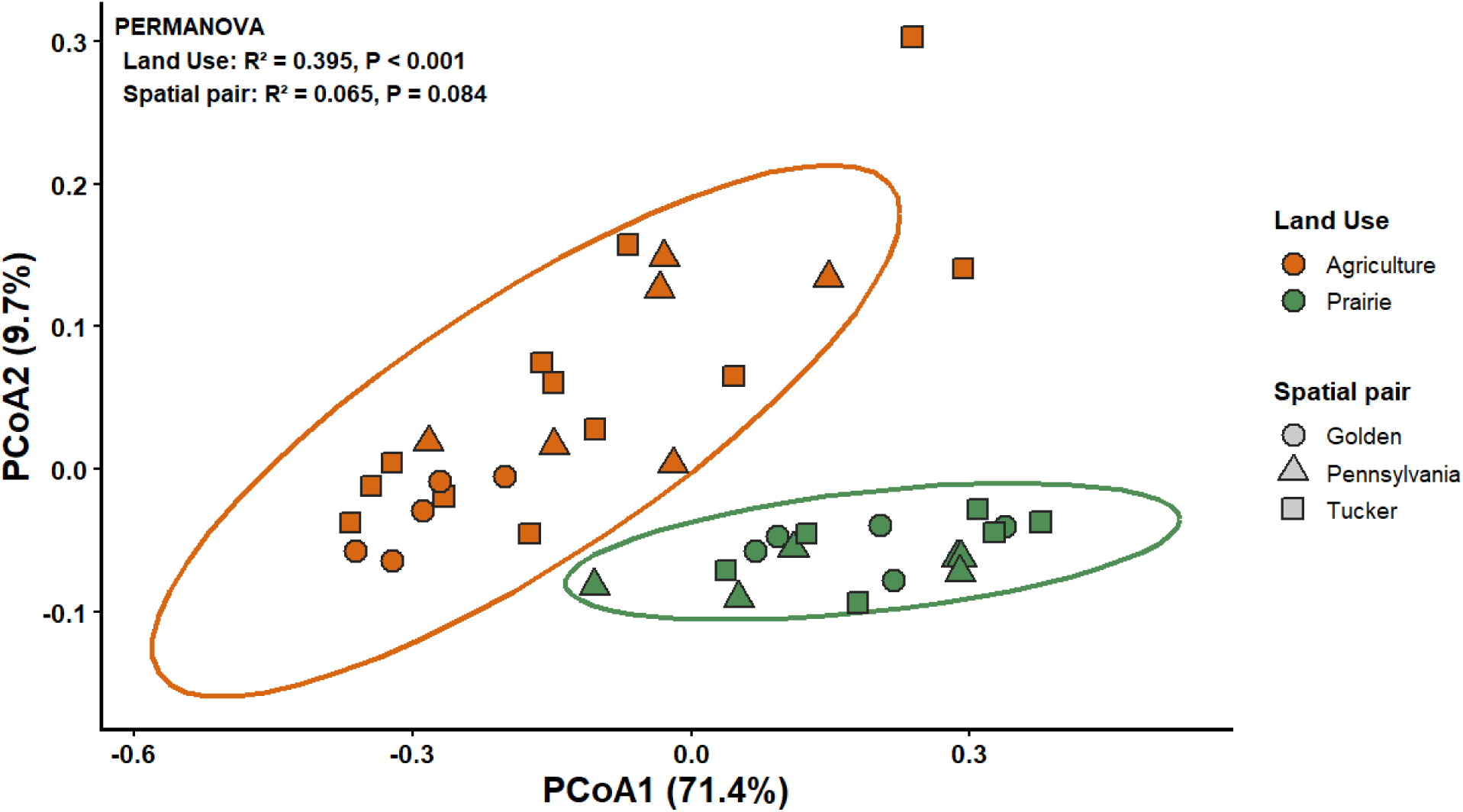
Principal coordinates analysis (PCoA) of Hellinger-transformed Ca. Udaeobacter recruitment profiles across prairie and agricultural soils. Recruitment counts were first converted to relative abundances within each sample and Hellinger transformed to remove differences in total transcript recruitment, allowing comparisons of genome composition among communities. Points represent individual metatranscriptomes, colors indicate land use (prairie or agriculture), and symbols denote spatial pairs (Golden, Penn, and Tucker). Ellipses represent the multivariate dispersion of each land use. The first two axes explained 71.4% and 9.7% of the total variation, respectively. Land use explained 39.5% of the variation in recruited genome composition after accounting for spatial pair (PERMANOVA, R² = 0.395, P < 0.001), whereas spatial pair explained only 6.5% of the variation and was not significant (R² = 0.065, P = 0.084). The clear separation of prairie and agricultural samples demonstrates that long-term land use restructures the composition of transcriptionally active Ca. Udaeobacter lineages independent of differences in overall recruitment abundance. **Alt text:** Principal coordinates analysis showing prairie and agricultural soil samples forming two separated clusters based on Hellinger-transformed Ca. Udaeobacter genome recruitment profiles, with ellipses indicating the spread of each land use group.

### Phylogenetic signal and genomic characteristics of land use-associated lineages

Land use-associated recruitment exhibited significant phylogenetic signal. Blomberg’s *K* indicated that closely related genomes showed more similar land use responses than expected by chance (*K* = 0.600, *P* = 0.040). Pagel’s lambda likewise supported strong phylogenetic structure (λ = 0.999, likelihood-ratio test, *P* < 0.0001), indicating that land use-associated recruitment closely followed the evolutionary relationships among the 36 consensus genomes. These results indicate that long-term land use did not influence individual Ca. Udaeobacter genomes independently, but land use-associated recruitment was concentrated within related evolutionary lineages (see Figure 3), consistently showing that the ecological traits underlying transcriptional recruitment are phylogenetically conserved.

**Figure 3.**
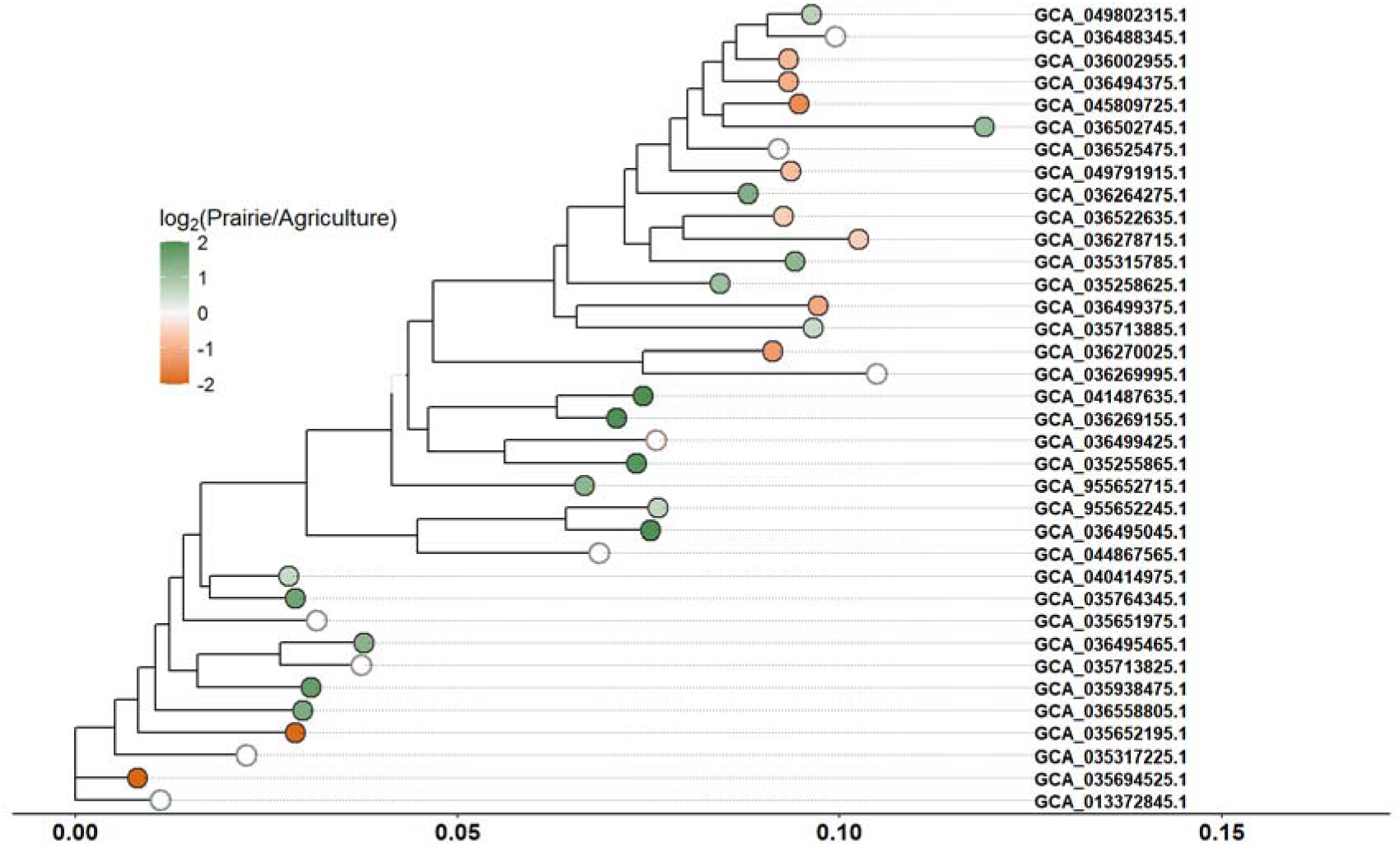
Phylogenetic distribution of land use-associated recruitment across the 36 consensus Ca. Udaeobacter genomes. The maximum-likelihood phylogeny is annotated by differential transcriptional recruitment between prairie and agricultural soils. Branch tip color indicates the DESeq2 log2 fold change, with green indicating greater recruitment in prairie soils and orange indicating greater recruitment in agricultural soils. Filled circles denote genomes that met the significance threshold (adjusted P < 0.05 and |log2 fold change| ≥ 0.5), whereas open circles represent genomes that did not meet these criteria. Land use-associated recruitment was strongly phylogenetically structured, with prairie-enriched genomes clustering within multiple related evolutionary lineages and comparatively fewer agricultural-enriched genomes distributed across the phylogeny, indicating that long-term land use preferentially altered transcriptional recruitment of evolutionarily conserved Ca. Udaeobacter lineages. **Alt text:** Maximum-likelihood phylogenetic tree of 36 Candidatus Udaeobacter genomes with branch tips colored by log2 fold change in land-use recruitment, showing prairie-enriched genomes clustered within related lineages and agriculture-enriched genomes distributed more broadly across the tree.

The phylogenetic recruitment was accompanied by differences in genome architecture. Prairie-enriched genomes were substantially smaller than agriculture-enriched genomes, with median genome sizes of 2.21 Mb and 3.29 Mb, respectively. Genome size was negatively correlated with land use response (ρ = −0.56, P = 0.002) (See Figure 4), indicating that progressively smaller genomes became increasingly associated with prairie recruitment. Despite their smaller genomes, prairie-associated lineages expressed a greater fraction of their predicted coding capacity than agriculture-associated lineages (34.7% versus 21.5%) and exhibited greater expressed functional richness (median 363 versus 264 expressed KOs). Genome size showed no relationship with assembly quality metrics or total recruitment, indicating that these differences were unlikely to reflect differences in assembly quality or sequencing depth. Combined, these results indicate that long-term land use not only restructures the phylogenetic composition of transcriptionally active Ca. Udaeobacter but also selects lineages with distinct genomic strategies. Prairie soils consistently favored smaller genomes that expressed a larger proportion of their coding capacity and a broader repertoire of functional genes, whereas larger genomes were more frequently associated with agricultural soils.

**Figure 4.**
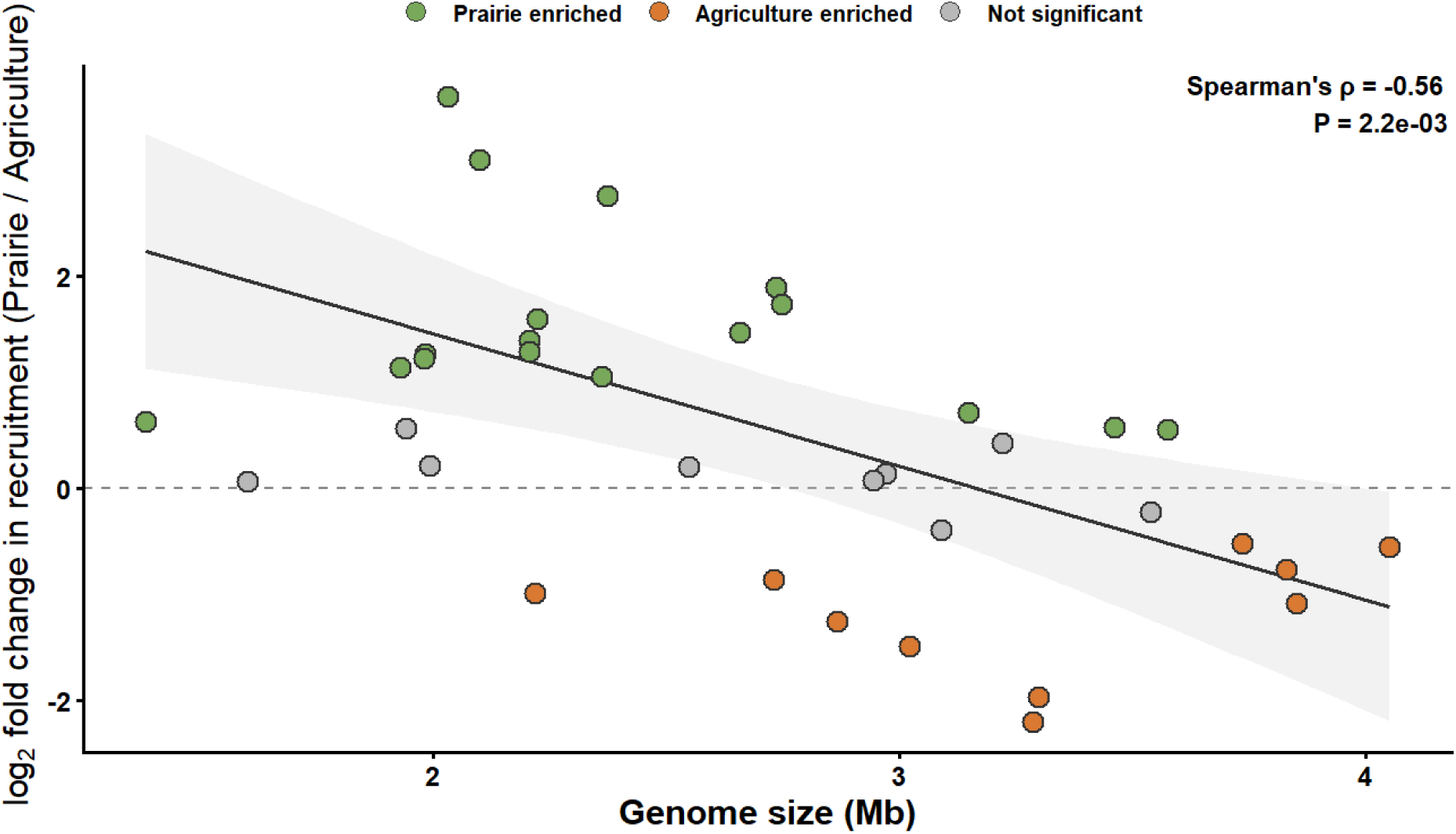
Genome size predicts land use-associated recruitment among transcriptionally active Ca. Udaeobacter lineages. Differential recruitment (DESeq2 log2 fold change, Prairie/Agriculture) is plotted against genome size for the 36 consensus Ca. Udaeobacter genomes. Genomes significantly enriched in prairie soils (adjusted P < 0.05, log2 fold change ≥ 0.5) are shown in green, genomes significantly enriched in agricultural soils (adjusted P < 0.05, log2 fold change ≤ −0.5) are shown in orange, and genomes not meeting these criteria are shown in gray. The fitted linear regression is shown for the land use-enriched genomes, with the shaded region representing the 95% confidence interval. Genome size was negatively associated with land use response (Spearman’s ρ = −0.56, P = 0.002), indicating that smaller genomes were preferentially recruited in native prairie soils, whereas larger genomes were more frequently associated with long-term agricultural management. **Alt text:** Scatter plot showing a negative relationship between genome size and land-use response, with a fitted regression line and shaded confidence band. Smaller genomes cluster toward prairie enrichment and larger genomes cluster toward agricultural enrichment.

## Discussion

By combining a curated, cross-referenced reference genome collection with metatranscriptomic recruitment, we demonstrate that long-term agricultural conversion reshapes not only the overall transcriptional activity of Ca. Udaeobacter, but also the evolutionary lineages that remain transcriptionally active. Rather than acting uniformly across the genus, land use selectively favors distinct phylogenetic groups, indicating that land use filtering operates at the lineage level within this globally abundant soil bacterium. These findings demonstrate that ecological responses within abundant soil microbial taxa are not homogeneous but instead reflect conserved evolutionary strategies that are shared across closely related lineages. Consequently, the effects of long-term land-use change extend beyond shifts in overall microbial activity to encompass the restructuring of evolutionary diversity within dominant soil bacterial groups. Ca. Udaeobacter represents one of the most widespread oligotrophic bacterial lineages in terrestrial ecosystems, so this work also establishes a framework for investigating how long-term environmental change influences other oligotrophic, carbon-conservative microorganisms. More broadly, integrating curated reference genome collections with metatranscriptomic recruitment provides a scalable approach for linking microbial activity with evolutionary trajectory, allowing ecological responses to be interpreted at the lineage level rather than solely through taxonomic abundance.

The normalized analyses demonstrate that these differences cannot be explained solely by the overwhelming reduction in total transcript recruitment. Even after standardizing recruitment within samples, prairie and agricultural communities remained compositionally distinct, indicating that long-term cultivation alters the identity of transcriptionally active lineages rather than simply suppressing activity across the genus [34]. This indicates that agricultural conversion functions as an ecological filter [35], replacing some lineages with others rather than uniformly suppressing activity across the genus. This level of lineage turnover indicates that closely related genomes occupy distinct ecological niches and differ in their ability to persist under contrasting soil environments created by perennial prairie vegetation versus annually manipulated agricultural systems [36]. This study shows the potential for long-term management to reshape the realized ecological niche of Ca. Udaeobacter by favoring evolutionary lineages possessing traits better suited to the altered physical, chemical, and biological conditions characteristic of agricultural soils.

Significant phylogenetic signal (Pagel’s λ and Blomberg’s K) indicates that land use-associated recruitment reflects conserved evolutionary traits rather than independent responses of individual genomes. A similar conceptual framework was proposed by Aguirre de Cárcer [37], discussing principles in community assembly upon selection pressures and the notion of phylogenetically conserved traits. Closely related genomes exhibited similar recruitment patterns across land-use types, showing that agricultural conversion does not influence Ca. Udaeobacter randomly across the phylogeny. Instead, long-term cultivation consistently favors some evolutionary lineages while greatly reducing the activity of others. This indicates that the traits allowing lineages to remain transcriptionally active under prairie or agricultural conditions have been retained through evolutionary history rather than arising independently in individual genomes [38,39]. The nearly eleven-fold reduction in overall transcriptional activity further indicates that agricultural soils do not simply shift activity among lineages but remove much of the evolutionary diversity that contributes to the active Ca. Udaeobacter community. The lineages found in this study that remain active in agricultural soils likely represent only a subset of the ecological and functional potential present in native prairie, reflecting the strength of environmental filtering imposed by long-term cultivation. The ecological consequences of this narrowing remain unknown, particularly whether the agricultural lineages retain the same functions expressed in prairie soils or occupy fundamentally different roles under long-term cultivation.

Genome size was one of the strongest predictors of land use-associated recruitment. Contrary to common expectations that long-term agricultural disturbance would favor smaller, streamlined genomes in some microbial consortia [6], the most strongly prairie-associated Ca. Udaeobacter lineages consistently possessed the smallest genomes while also expressing a larger proportion of their predicted coding sequences. These findings reveal that genome reduction in *Ca.* Udaeobacter is not simply a strategy for surviving resource-poor environments, a description that is itself a misnomer for native tallgrass prairie soils [40]. Instead, smaller genomes appear to be associated with ecological specialization for stable perennial grassland soils, where efficient resource acquisition and sustained interactions with surrounding microbial communities may provide a greater advantage than maintaining broad metabolic flexibility [41]. In contrast, the lineages that remain transcriptionally active after decades of agricultural management tended to possess larger genomes despite the dramatic reduction in overall activity observed across the genus. This pattern implies that persistence under long-term cultivation may depend less on maximizing transcriptional output and more on maintaining a broader suite of physiological capabilities that allow survival under the more variable physical, chemical, and biological conditions characteristic of agricultural soils. In addition, the energetic cost of larger genomes may be favored under variable, yet high energy input conditions that come with agricultural production; albeit often yielding less carbon substrates while increasing inorganic forms of nutrients [42]. Together with the strong phylogenetic signal observed across consensus phylogeny, these results show both genome architecture and land use preference are evolutionarily conserved within Ca. Udaeobacter. Long-term land use therefore acts as an ecological filter operating below the genus level, preferentially retaining evolutionary lineages that share conserved genomic strategies while excluding others from the transcriptionally active community [43]. Given the century-long timescales represented by these sites, this filtering reflects more than a temporary ecological response. Persistent environmental selection pressure has altered which evolutionary lineages actively contribute to the soil microbiome, potentially shifting the evolutionary trajectory of the active Ca. Udaeobacter community [44].

Because Ca. Udaeobacter is among the most abundant bacterial groups in soils worldwide [45], this filtering likely extends beyond changes in taxonomic composition to a loss of evolutionary and functional diversity. Rather than simply reducing the activity of a dominant soil bacterium, agricultural conversion appears to narrow the range of ecological strategies contributing to the active microbiome, leaving a smaller subset of lineages capable of maintaining transcription under managed conditions. If lineages lost from agricultural soils are not readily maintained or reestablished, restoration may require rebuilding evolutionary diversity rather than simply recovering microbial abundance or activity.

Several limitations should be considered when interpreting these findings. First, this study focused on paired prairie and agricultural sites within Missouri, and broader geographic sampling is necessary to determine whether similar recruitment patterns occur across the wider distribution of North American grasslands and agricultural systems. Second, recruitment was evaluated relative to a curated consensus reference composed of currently available Candidatus Udaeobacter genomes. Although this approach substantially improved taxonomic confidence by requiring agreement between NCBI and GTDB classifications, additional uncultivated diversity will undoubtedly become available as new genomes are recovered. Finally, transcript recruitment reflects the activity of reference-associated populations rather than absolute organism abundance. Future studies integrating metagenomics, metatranscriptomics, or MAG-based functional characterization will help determine the specific physiological traits responsible for the strong phylogenetic patterns observed here.

Long-term agricultural management has reshaped Ca. Udaeobacter at multiple levels of biological organization. Beyond reducing overall transcriptional activity, it filters evolutionary lineages, favors distinct genomic strategies, and narrows the diversity of lineages contributing to the active soil microbiome. These results demonstrate that the legacy of cultivation extends beyond short-term ecological responses and into the evolutionary composition of one of the most abundant soil bacterial groups, encompassing changes in the evolutionary composition of transcriptionally active soil microbial communities more broadly. This study demonstrates that combining curated reference genome collections with metatranscriptomic recruitment provides a powerful framework for resolving lineage-level ecological responses within abundant but poorly understood microbial taxa, offering a new approach for linking microbial evolution with ecosystem function.

## Supporting information

Supplemental Table 1. Assembly Statistics

## Data Availability

The metatranscriptomic sequencing data are part of a larger dataset supporting ongoing, unpublished research on separate questions, and are not yet publicly available in order to protect the priority of that work. Data will be deposited in the NCBI Sequence Read Archive upon publication of the related studies, and are available from the corresponding author upon reasonable request in the interim. The 36 *Candidatus* Udaeobacter consensus reference genomes used in this study are previously published, publicly available assemblies; their NCBI accessions are provided in Table S1.

## Acknowledgements

This research was supported by the USDA-ARS under agreement No. 59-6020-5-001, and the University of Missouri Center for Agroforestry and the USDA-ARS Dale Bumpers Small Farm Research Center.

---

**Table S1.** Assembly statistics for the 36 taxonomically concordant *Candidatus* Udaeobacter consensus reference genomes. Genome size, contig count, largest contig length, N50, and GC content are reported for each NCBI accession retained after cross-referencing against GTDB (release r232) taxonomic assignments. These genomes constitute the consensus reference collection used for phylogenomic reconstruction and transcript recruitment throughout this study.

| Accession | Genome size (Mb) | Contigs | Largest contig (kb) | N50 (kb) | GC (%) |
| --- | --- | --- | --- | --- | --- |
| GCA_013372845.1 | 3.222 | 145 | 92.97 | 24.81 | 55.16 |
| GCA_035255865.1 | 2.735 | 267 | 71.22 | 14.19 | 55.27 |
| GCA_035258625.1 | 2.363 | 278 | 42.28 | 10.17 | 55.09 |
| GCA_035315785.1 | 1.981 | 164 | 67.08 | 17.58 | 54.26 |
| GCA_035317225.1 | 2.549 | 306 | 73.3 | 11.95 | 54.3 |
| GCA_035651975.1 | 2.97 | 71 | 234.42 | 70.72 | 54.68 |
| GCA_035652195.1 | 3.298 | 172 | 121.25 | 31.13 | 53.9 |
| GCA_035694525.1 | 3.287 | 441 | 63.74 | 12.05 | 54.55 |
| GCA_035713825.1 | 3.538 | 205 | 134.7 | 29.83 | 53.9 |
| GCA_035713885.1 | 3.576 | 204 | 104.68 | 24.32 | 53.73 |
| GCA_035764345.1 | 2.224 | 449 | 24.59 | 5.91 | 54.6 |
| GCA_035938475.1 | 2.748 | 154 | 98.51 | 32.32 | 54.26 |
| GCA_036002955.1 | 2.73 | 304 | 38 | 11.61 | 54.46 |
| GCA_036264275.1 | 2.207 | 299 | 37.27 | 9.52 | 54.28 |
| GCA_036269155.1 | 2.032 | 287 | 37.01 | 7.99 | 54.78 |
| GCA_036269995.1 | 1.604 | 51 | 123.96 | 42.52 | 54.94 |
| GCA_036270025.1 | 2.866 | 478 | 40.57 | 8.55 | 55.87 |
| GCA_036278715.1 | 3.735 | 106 | 169.56 | 55.78 | 54.76 |
| GCA_036488345.1 | 2.944 | 231 | 138.33 | 20.4 | 54.79 |
| GCA_036494375.1 | 2.22 | 368 | 36.45 | 8.58 | 54.69 |
| GCA_036495045.1 | 2.374 | 143 | 238.69 | 30.76 | 54.47 |
| GCA_036495465.1 | 2.208 | 190 | 60.37 | 15.66 | 53.65 |
| GCA_036499375.1 | 3.851 | 67 | 314.6 | 97.09 | 55.12 |
| GCA_036499425.1 | 3.091 | 153 | 128.09 | 37.51 | 55.06 |
| GCA_036502745.1 | 1.931 | 336 | 41.97 | 6.35 | 54.36 |
| GCA_036522635.1 | 4.051 | 283 | 63.56 | 22.64 | 54.16 |
| GCA_036525475.1 | 1.944 | 411 | 15.99 | 5.12 | 54.97 |
| GCA_036558805.1 | 2.658 | 248 | 60.57 | 12.9 | 54.09 |
| GCA_040414975.1 | 3.461 | 419 | 39.52 | 11.39 | 54.23 |
| GCA_041487635.1 | 2.101 | 327 | 36.46 | 7.48 | 54.9 |
| GCA_044867565.1 | 1.993 | 292 | 31.5 | 8.07 | 55.26 |
| GCA_045809725.1 | 3.022 | 259 | 61.6 | 16.67 | 54.74 |
| GCA_049791915.1 | 3.83 | 288 | 123.4 | 23.4 | 55.22 |
| GCA_049802315.1 | 3.149 | 589 | 51.41 | 6.6 | 54.89 |
| GCA_955652245.1 | 1.385 | 210 | 24.13 | 7.78 | 54.71 |
| GCA_955652715.1 | 1.984 | 369 | 22.59 | 5.67 | 55.03 |

## References

[1] Sampson F, Knopf F. Prairie conservation in North America. BioScience 1994;44:418–421. 10.2307/1312365

[2] Henderson DC, Koper N. Historic distribution and ecology of tall-grass prairie in western Canada; 2014.

[3] Hartman K, Van Der Heijden MGA, Wittwer RA, et al. Cropping practices manipulate abundance patterns of root and soil microbiome members paving the way to smart farming. Microbiome 2018;6:14. 10.1186/s40168-017-0389-9

[4] Naziębło A, Pytlak A, Furtak A, et al. Advances and hotspots in research on Verrucomicrobiota: focus on agroecosystems. Microb Ecol 2026;89:1. 10.1007/s00248-025-02657-3

[5] Klopf RP, Baer SG, Bach EM, et al. Restoration and management for plant diversity enhances the rate of belowground ecosystem recovery. Ecol Appl 2017;27:355–362. 10.1002/eap.1503

[6] Simonsen AK. Environmental stress leads to genome streamlining in a widely distributed species of soil bacteria. ISME J 2022;16:423–434. 10.1038/s41396-021-01082-x

[7] Hassan-Dalléac S, Guiga W, Suau-Pernet A. Soil microbes are the tiny bioengineers running Earth’s underground factory. Commun Earth Environ 2026;7:403. 10.1038/s43247-026-03544-6

[8] Bergmann GT, Bates ST, Eilers KG, et al. The under-recognized dominance of Verrucomicrobia in soil bacterial communities. Soil Biol Biochem 2011;43:1450–1455. 10.1016/j.soilbio.2011.03.012

[9] Brewer TE, Handley KM, Carini P, et al. Genome reduction in an abundant and ubiquitous soil bacterium ‘Candidatus Udaeobacter copiosus’. Nat Microbiol 2016;2:16198. 10.1038/nmicrobiol.2016.198

[10] Willms IM, Rudolph AY, Göschel I, et al. Globally abundant ‘Candidatus Udaeobacter’ benefits from release of antibiotics in soil and potentially performs trace gas scavenging. mSphere 2020;5:e00186–20. 10.1128/mSphere.00186-20

[11] Wattenburger CJ, Buckley DH. Land use alters bacterial growth dynamics in soil. Environ Microbiol 2023;25:3239–3254. 10.1111/1462-2920.16514

[12] Hu J, Richwine JD, Keyser PD, et al. Nitrogen fertilization and native C4 grass species alter abundance, activity, and diversity of soil diazotrophic communities. Front Microbiol 2021;12:675693. 10.3389/fmicb.2021.675693

[13] Semchenko M, Barry KE, Vries FT, et al. Deciphering the role of specialist and generalist plant–microbial interactions as drivers of plant–soil feedback. New Phytol 2022;234:1929– 1944. 10.1111/nph.18118

[14] Suleiman AKA, Manoeli L, Boldo JT, et al. Shifts in soil bacterial community after eight years of land-use change. Syst Appl Microbiol 2013;36:137–144. 10.1016/j.syapm.2012.10.007

[15] Montecchia MS, Tosi M, Soria MA, et al. Pyrosequencing reveals changes in soil bacterial communities after conversion of Yungas forests to agriculture. PLoS One 2015;10:e0119426. 10.1371/journal.pone.0119426

[16] Kitts PA, Church DM, Thibaud-Nissen F, et al. Assembly: a resource for assembled genomes at NCBI. Nucleic Acids Res 2016;44:D73–D80. 10.1093/nar/gkv1226

[17] Parks DH, Chuvochina M, Rinke C, et al. GTDB: an ongoing census of bacterial and archaeal diversity through a phylogenetically consistent, rank normalized and complete genome-based taxonomy. Nucleic Acids Res 2022;50:D785–D794. 10.1093/nar/gkab776

[18] Langmead B, Salzberg SL. Fast gapped-read alignment with Bowtie 2. Nat Methods 2012;9:357–359. 10.1038/nmeth.1923

[19] Li H, Handsaker B, Wysoker A, et al. The Sequence Alignment/Map format and SAMtools. Bioinformatics 2009;25:2078–2079. 10.1093/bioinformatics/btp352

[20] Aramaki T, Blanc-Mathieu R, Endo H, et al. KofamKOALA: KEGG Ortholog assignment based on profile HMM and adaptive score threshold. Bioinformatics 2020;36:2251–2252. 10.1093/bioinformatics/btz859

[21] Chaumeil P-A, Mussig AJ, Hugenholtz P, et al. GTDB-Tk v2: memory friendly classification with the genome taxonomy database. Bioinformatics 2022;38:5315–5316. 10.1093/bioinformatics/btac672

[22] Wong TKF, Ly-Trong N, Ren H, et al. IQ-TREE 3: phylogenomic inference software using complex evolutionary models. Mol Biol Evol 2026;43:msag117. 10.1093/molbev/msag117

[23] R Core Team. R: a language and environment for statistical computing. Vienna, Austria: R Foundation for Statistical Computing; 2026. Available at: https://www.R-project.org/

[24] Love MI, Huber W, Anders S. Moderated estimation of fold change and dispersion for RNA-seq data with DESeq2. Genome Biol 2014;15:550. 10.1186/s13059-014-0550-8

[25] Benjamini Y, Hochberg Y. Controlling the false discovery rate: a practical and powerful approach to multiple testing. J R Stat Soc Series B Stat Methodol 1995;57:289–300. 10.1111/j.2517-6161.1995.tb02031.x

[26] Legendre P, Gallagher ED. Ecologically meaningful transformations for ordination of species data. Oecologia 2001;129:271–280. 10.1007/s004420100716

[27] Oksanen J, Simpson G, Blanchet F, et al. vegan: community ecology package. R package version 2.8-0; 2026. https://vegandevs.github.io/vegan/

[28] Anderson MJ. Distance-based tests for homogeneity of multivariate dispersions. Biometrics 2006;62:245–253. 10.1111/j.1541-0420.2005.00440.x

[29] Blomberg SP, Garland T, Ives AR. Testing for phylogenetic signal in comparative data: behavioral traits are more labile. Evolution 2003;57:717–745. 10.1111/j.0014-3820.2003.tb00285.x

[30] Mitteroecker P, Collyer ML, Adams DC. Exploring phylogenetic signal in multivariate phenotypes by maximizing Blomberg’s K. Syst Biol 2025;74:215–229. 10.1093/sysbio/syae035

[31] Pearse WD, Davies TJ, Wolkovich EM. How to define, use, and interpret Pagel’s λ (lambda) in ecology and evolution. Glob Ecol Biogeogr 2025;34:e70012. 10.1111/geb.70012

[32] Pagel M. Inferring the historical patterns of biological evolution. Nature 1999;401:877–884. 10.1038/44766

[33] Revell LJ. phytools: an R package for phylogenetic comparative biology (and other things). Methods Ecol Evol 2012;3:217–223. 10.1111/j.2041-210X.2011.00169.x

[34] Tibbett M, Fraser TD, Duddigan S. Identifying potential threats to soil biodiversity. PeerJ 2020;8:e9271. 10.7717/peerj.9271

[35] Aguilar-Trigueros CA, Rillig MC, Ballhausen M-B. Environmental filtering is a relic. A response to Cadotte and Tucker. Trends Ecol Evol 2017;32:882–884. 10.1016/j.tree.2017.09.013

[36] Goberna M, García C, Verdú M. A role for biotic filtering in driving phylogenetic clustering in soil bacterial communities. Glob Ecol Biogeogr 2014;23:1346–1355. 10.1111/geb.12227

[37] Aguirre de Cárcer D. A conceptual framework for the phylogenetically constrained assembly of microbial communities. Microbiome 2019;7:142. 10.1186/s40168-019-0754-y

[38] Hargreaves SK, Williams RJ, Hofmockel KS. Environmental filtering of microbial communities in agricultural soil shifts with crop growth. PLoS One 2015;10:e0134345. 10.1371/journal.pone.0134345

[39] Causevic S, Tackmann J, Sentchilo V, et al. Habitat filtering more than microbiota origin controls microbiome transplant outcomes in soil. ISME J 2025;19:wraf162. 10.1093/ismejo/wraf162

[40] Chuckran PF, Hungate BA, Schwartz E, et al. Variation in genomic traits of microbial communities among ecosystems. FEMS Microbes 2022;2:xtab020. 10.1093/femsmc/xtab020

[41] Goodall T, Busi SB, Griffiths RI, et al. Soil properties in agricultural systems affect microbial genomic traits. FEMS Microbes 2025;6:xtaf008. 10.1093/femsmc/xtaf008

[42] Piton G, Allison SD, Bahram M, et al. Life history strategies of soil bacterial communities across global terrestrial biomes. Nat Microbiol 2023;8:2093–2102. 10.1038/s41564-023-01465-0

[43] Talavera-Marcos S, Parras-Moltó M, Aguirre de Cárcer D. Leveraging phylogenetic signal to unravel microbiome function and assembly rules. Comput Struct Biotechnol J 2023;21:5165–5173. 10.1016/j.csbj.2023.10.039

[44] Wagg C, Hautier Y, Pellkofer S, et al. Diversity and asynchrony in soil microbial communities stabilizes ecosystem functioning. eLife 2021;10:e62813. 10.7554/eLife.62813

[45] Willms IM, Bolz SH, Yuan J, et al. The ubiquitous soil verrucomicrobial clade ‘Candidatus Udaeobacter’ shows preferences for acidic pH. Environ Microbiol Rep 2021;13:878–883. 10.1111/1758-2229.13006

